# cAMP-Fyn signaling in the dorsomedial striatum direct pathway drives excessive alcohol use

**DOI:** 10.1101/2020.01.07.898023

**Authors:** Yann Ehinger, Nadege Morisot, Khanhky Phamluong, Samuel A. Sakhai, Drishti Soneja, Martin F. Adrover, Veronica A. Alvarez, Dorit Ron

## Abstract

Fyn kinase in the dorsomedial striatum (DMS) of rodents plays a central role in mechanisms underlying excessive alcohol intake. The DMS is comprised of medium spiny neurons (MSNs) that project directly (dMSNs) or indirectly (iMSNs) to the substantia nigra. Here, we examined the cell-type specificity of Fyn’s actions on alcohol use. First, we knocked down Fyn selectively in DMS dMSNs or iMSNs of mice and measured the level of alcohol consumption. We found that downregulation of Fyn in dMSNs, but not in iMSNs, reduces excessive alcohol but not saccharin intake. D1Rs are coupled to Gαs/olf, which activate cAMP signaling. To examine whether Fyn’s actions are mediated through cAMP signaling, DMS dMSNs were infected with GαsDREADD, and the activation of Fyn signaling was measured following CNO treatment. We found that remote stimulation of cAMP signaling in DMS dMSNs activates Fyn and promotes the phosphorylation of the Fyn substrate, GluN2B. In contract, remote activation of GαsDREADD in DLS dMSNs did not alter Fyn signaling. We then tested whether activation of GαsDREADD in DMS dMSNs or iMSNs alters alcohol intake and observed that CNO-dependent activation of GαsDREADD in DMS dMSNs but not iMSNs increases alcohol but not saccharin intake. Finally, we examined the contribution of Fyn to GαsDREADD-dependent increase in alcohol intake, and found that systemic administration of the Fyn inhibitor, AZD0503 blocks GαsDREADD-dependent increase in alcohol consumption. Our results suggest that the cAMP-Fyn axis in the DMS dMSNs is a molecular transducer of mechanisms underlying the development of excessive alcohol consumption.

## Introduction

The dorsomedial striatum (DMS) is critically involved in processes such as locomotion [1], and goal-directed behaviors [2,3]. The DMS is comprised primarily of GABAergic medium spiny projection neurons (MSNs) that receive dopaminergic input from the midbrain [4]. MSNs can be divided into two populations of neurons that take part in opposing activities [5]; MSNs that project directly to the substantia nigra pars reticula (SNr) facilitate actions and are defined as direct MSNs (dMSNs) [5], and MSNs that project indirectly to the SNr gate actions and are defined as indirect MSNs (iMSNs) [5]. dMSNs selectively express the dopamine D1 receptors (D1Rs) whereas iMSNs express the dopamine D2 receptors (D2Rs) [5]. In the striatum, D1Rs are coupled to Gαolf, a homolog of Gαs [6]. Stimulation of Gαs/olf-coupled receptors results in the production of the second messenger cyclic adenosine monophosphate (cAMP) [6,7], which binds to, and activates, protein kinase A (PKA) [8], a kinase that plays an important role in the adult brain [9-11]. In contrast, D2Rs are coupled to Gαi, which inhibits cAMP signaling [7]. dMSNs and iMSNs exert balanced influence on locomotion and goal-directed behaviors [5], and an imbalance of dMSNs and iMSNs function has been implicated in neurodegenerative disorders such as Parkinson’s disease [5,12], as well as psychiatric disorders such as obsessive compulsive disorder, anxiety and addiction [13-15].

We previously observed that Fyn kinase is activated in DMS dMSNs upon stimulation of D1Rs [16]. Fyn belongs to the Src family of non-receptor protein tyrosine kinases (PTKs) [17,18], and is highly expressed in the developing and adult brain in regions such as cortex, hippocampus and cerebellum as well as in the striatum [19,20]. Fyn plays an important role in the CNS [21], as it modulates excitatory and inhibitory synaptic transmission and participates in learning and memory processes [21-28]. Dysfunction of Fyn signaling has been associated with Alzheimer’s disease [29] and pain [23].

Accumulating data in humans and rodents also suggest that Fyn plays a central role in cellular neuroadaptations that underlie alcohol use disorder (AUD) [30,31]. Specifically, genetic mutations within the Fyn gene have been associated with increased susceptibility for the development and severity of AUD in humans [32-34], and gene network association studies identified a link between Fyn and alcohol dependence [35]. Animal data suggest that Fyn plays a role in the acute tolerance to the hypnotic sedative effect of alcohol [36,37], as well as in alcohol drinking behavior [38-40]. Molecularly, excessive consumption of alcohol activates Fyn specifically in the DMS of mice and rats [39,41,42]. Once activated by alcohol, Fyn phosphorylates its substrate, GluN2B [39,41,42]. Alcohol-dependent Fyn-mediated phosphorylation of GluN2B produces in forward trafficking of the channel and long-lasting enhancement of GluN2B activity in the DMS [39]. Inhibition of Fyn in the DMS of rats attenuates operant self-administration of alcohol [39], and systemic administration of the Fyn inhibitor, AZD0530, attenuates goal-directed alcohol seeking and facilitates extinction in mice [40]. Together, these data suggest that Fyn in the DMS plays a central role in neuroadaptations that underlie alcohol use.

This study was aimed to explore the cellular specificity of Fyn-dependent molecular and behavioral neuroadaptations that drive AUD.

## Methods

The description of purchased reagents, collection of brain samples, western blot analysis, immunoprecipitation, preparation of solutions and the preparation of FLEX-shRNA-Fyn and FLEX-SCR is detailed in the Supplementary Information section.

### Animals

C57BL/6 mice obtained from Jackson Laboratories. Drd1a-Cre (D1-Cre) and AdoraA2-Cre (A2A-Cre) mice both of which are on C57BL/6 background, were obtained from Mutant Mice Resource and Research Centers (MMRRC) UC Davis (David, CA). Ai14 mice were purchased from Jackson Laboratory (Bar Harbor, Maine). The generation of D1-Cre/Ai14 mouse line is described in (Wang et al, 2015). The same breeding strategy was used to generate the A2A-Cre/Ai14 mouse line. Mice were genotyped by polymerase chain reaction (PCR) analysis of products derived from tail DNA. Male mice were 8-9 weeks old at the beginning of the experiments and were individually housed in temperature and humidity-controlled rooms under a reversed 12-hours light/dark cycle. Food and water were available *ad libitum*. All animal procedures were approved by the University of California San Francisco (UCSF) Institutional Animal Care and Use Committee and were conducted in agreement with the Association for Assessment and Accreditation of Laboratory Animal Care (AAALAC, UCSF).

### Infection of the DMS with FLEX-shFyn and FLEX-SCR

D1-Cre/Ai14 and A2A-Cre/Ai14 mice were anesthetized using a mixture of ketamine (120 mg/kg) and xylazine (8 mg/kg). Bilateral microinfusions were made using stainless steel injectors (33 gauge, Small Parts) into the DMS (the stereotaxic coordinates were anterioposterior +1.1 mm from bregma; mediolateral ±1.2 mm from bregma and dorsoventral −2.8 and from bregma for the first injection site and anterioposterior +1.3 mm from bregma; mediolateral ±1.2 mm from bregma and dorsoventral −3 mm from bregma for the second injection site). Animals were infused with lenti-virus expressing FLEX-shFyn or its scramble control (FLEX-SCR) (1.2 μl/site with 2 sites of injection per hemisphere) at a concentration of 10^7^ pg/ml at an injection rate of 0.1 μl/minute [16]. After each infusion, the injectors were left in place for an additional 10 minutes to allow the virus to diffuse.

### Infection of the DMS and the dorsolateral striatum (DLS) with AAV-DIO-rM3D(Gs)-mCherry

D1-Cre or A2A-Cre mice were anesthetized using a mixture of ketamine (120 mg/kg) and xylazine (8 mg/kg). Bilateral microinfusions were made using stainless steel injectors (33 gauge, Small Parts) into the DMS (the stereotaxic coordinates were anterioposterior +1.1 mm from bregma; mediolateral ±1.25 mm from bregma and dorsoventral −2.8 mm from bregma) or the DLS (the stereotaxic coordinates were anterioposterior +1.1, medialateral ± 2.3 from bregma and dorsoventral −2.8 from bregma). Mice were infused with AAV-DIO-rM3D(Gs)-mCherry (AAV-DIO-Gs-DREADD) (1 μl per hemisphere) at a concentration of 10^13^ vg/ml and at an injection rate of 0.1 μl/minute. After each infusion, the injectors were left in place for an additional 10 minutes to allow the virus to diffuse.

### Drinking Paradigm

#### Two bottle choice - 20% alcohol

D1-Cre/Ai14 and A2A-Cre/Ai14 mice underwent one week of 2-bottle choice 20% (v/v) alcohol drinking paradigm (IA20%2BC) as described in [43,44]. Specifically, one month after stereotaxic surgery and infection of the DMS with FLEX-shFyn or FLEX-SCR, mice had 24 hours access to one bottle of 20% alcohol and one bottle of water on Monday, Wednesday and Friday with alcohol drinking sessions starting 2 hours into the dark cycle. During the 24 or 48 hours (weekend) of alcohol deprivation periods, mice had access to two bottles of water (**Figure 1b**).

**Figure 1.**
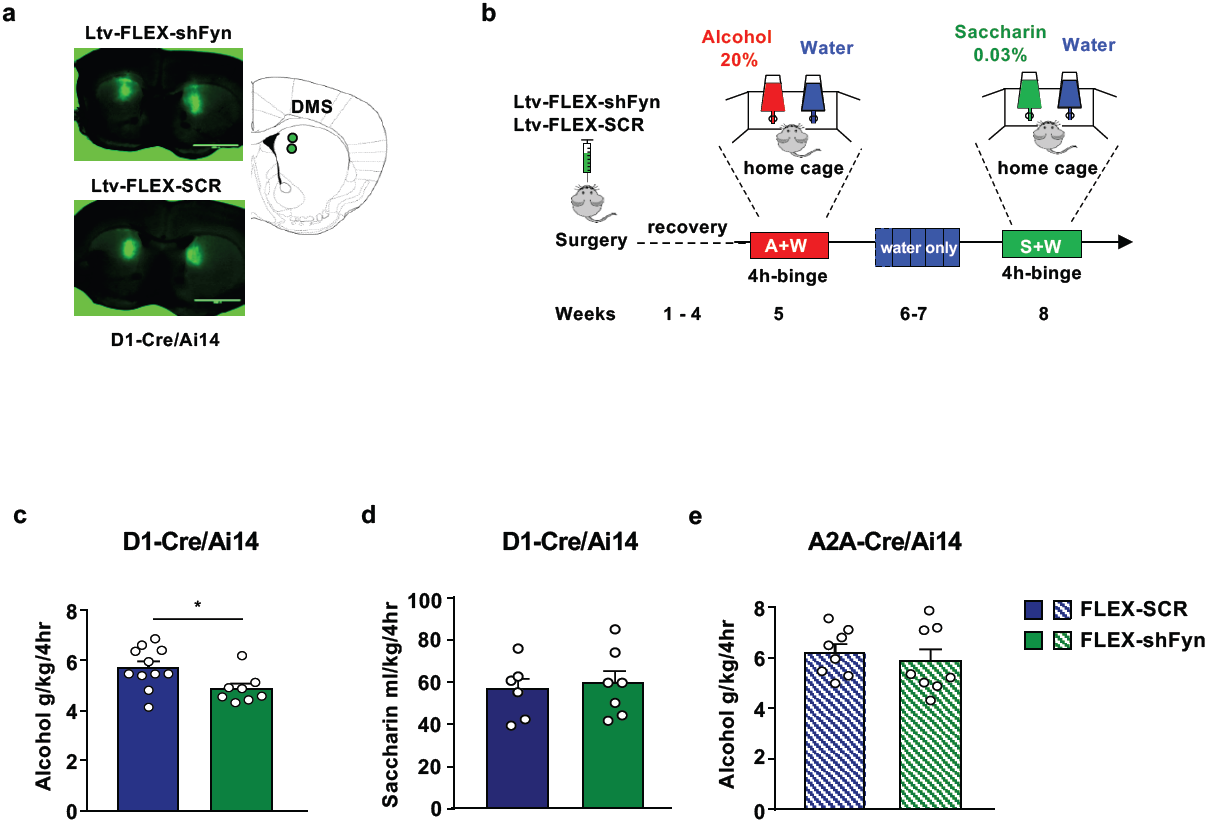
Knockdown of Fyn in DMS dMSNs but not in iMSNs attenuates the development of alcohol but not saccharin intake. **(a)** The DMS of D1-Cre/Ai14 mice was infected bilaterally with a lenti-virus expressing FLEX-shFyn or FLEX-SCR (10^7^pg/ml, 1.2 μl per site, 2 injection per side), and viral infection was evaluated 4 weeks later. Infection was visualized on EVOS FL microscope (Life Technologies, Carlsbad, CA), scale: 2x. (**b**) Timeline of experiments. Four weeks after surgery, D1-Cre/Ai14 (**c**) or A2A-Cre/Ai14 mice (**e**) underwent 20%2BC 4h binge drinking session, and alcohol and water intake were recorded. (**d**) At the completion of the alcohol drinking regimen, D1-Cre/Ai14 mice had access to water only for two weeks, followed by one week of home cage intermittent access to 0.03% saccharin. Saccharin and water intake were measured at the 4-hour time point. (**c**) Knockdown of Fyn in the DMS of D1-Cre/Ai14 mice attenuates alcohol intake (two-tailed unpaired t-test, t=2.491, p=0.0234). (**d**) Knockdown of Fyn in the DMS of D1-Cre/Ai14 mice does not alter saccharin intake (two-tailed unpaired t-test, t=0.2163, p=0.8327). (**e**) Knockdown of Fyn in the DMS of A2A-Cre/Ai14 does not affect alcohol intake (two-tailed unpaired t-test, t=0.583, p=0.5692). Data are presented as individual data points and mean ± SEM. * p<0.05. (**c**) FLEX-shFyn n=8, FLEX-SCR n=11. **(d**) FLEX-shFyn n=7, FLEX-SCR n=6, (**e**) FLEX-shFyn n=8, FLEX-SCR n=8.

#### Two bottle choice - 10% alcohol

One month after stereotaxic surgery and the infection of the DMS of D1-Cre or D2-Cre with AAV-DIO-Gs-DREADD, mice were habituated by intraperitoneal (IP) injection of saline for three days. On test day, mice received a systemic administration of vehicle (0.5% DMSO) or Clozapine N-Oxide (CNO, 3 mg/kg). Fifteen minutes later, mice had access to one bottle of 10% (v/v) alcohol and one bottle of water. Alcohol and water intake were measured 4 hours later (**Timeline, Figure 4a**). Mice were then given water only for one week and were tested again using a counterbalanced, within-subject design.

**Figure 2.**
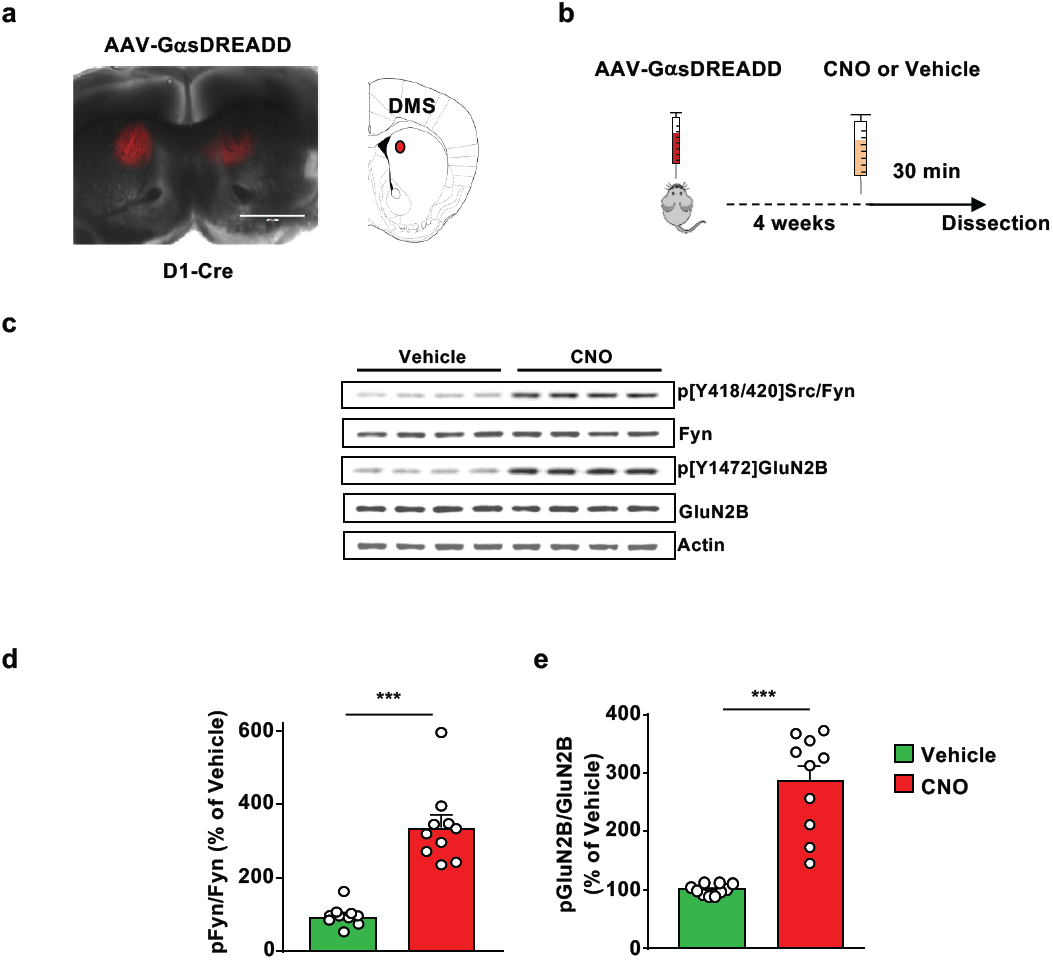
Stimulation of GαsDREADD in DMS dMSNs activates Fyn/GluN2B signaling. **(a)** The DMS of D1-Cre mice was infected bilaterally with AAV-DIO-rM3D(Gs)-mCherry (10^13^ vg/ml, 1 μl per side), and infection was evaluated 4 weeks later. Infection was visualized on EVOS FL microscope (Life Technologies, Carlsbad, CA), scale: 2x. (**b**) Timeline of experiment. Four weeks after surgery, vehicle (0.5% DMSO) or CNO (3 mg/kg) was systemically administered, and the DMS and was dissected 30 minutes later. (**c**) Representative image of Fyn activation and GluN2B phosphorylation which were measure by Western blot analysis using anti-Tyr417/420[Src/Fyn] and anti-Tyr1472[GluN2B] antibodies, respectively. Total protein levels of Fyn, GluN2B and actin, which was used as a loading control, were measured in parallel. (**d-e**) Data are presented as the individual data points and mean densitometry values of the phosphorylated protein divided by the densitometry values of the total protein ± SEM and expressed as % of vehicle. Activation of GαsDREADD in DMS dMSNs increases Fyn activation (**d**) (two-tailed unpaired t-test, t=7.148, p<0.001, and GluN2B phosphorylation (**e**) (two-tailed unpaired t-test, t=6.948, p<0.001). n=10 per treatment. ***p<0.001.

**Figure 3.**
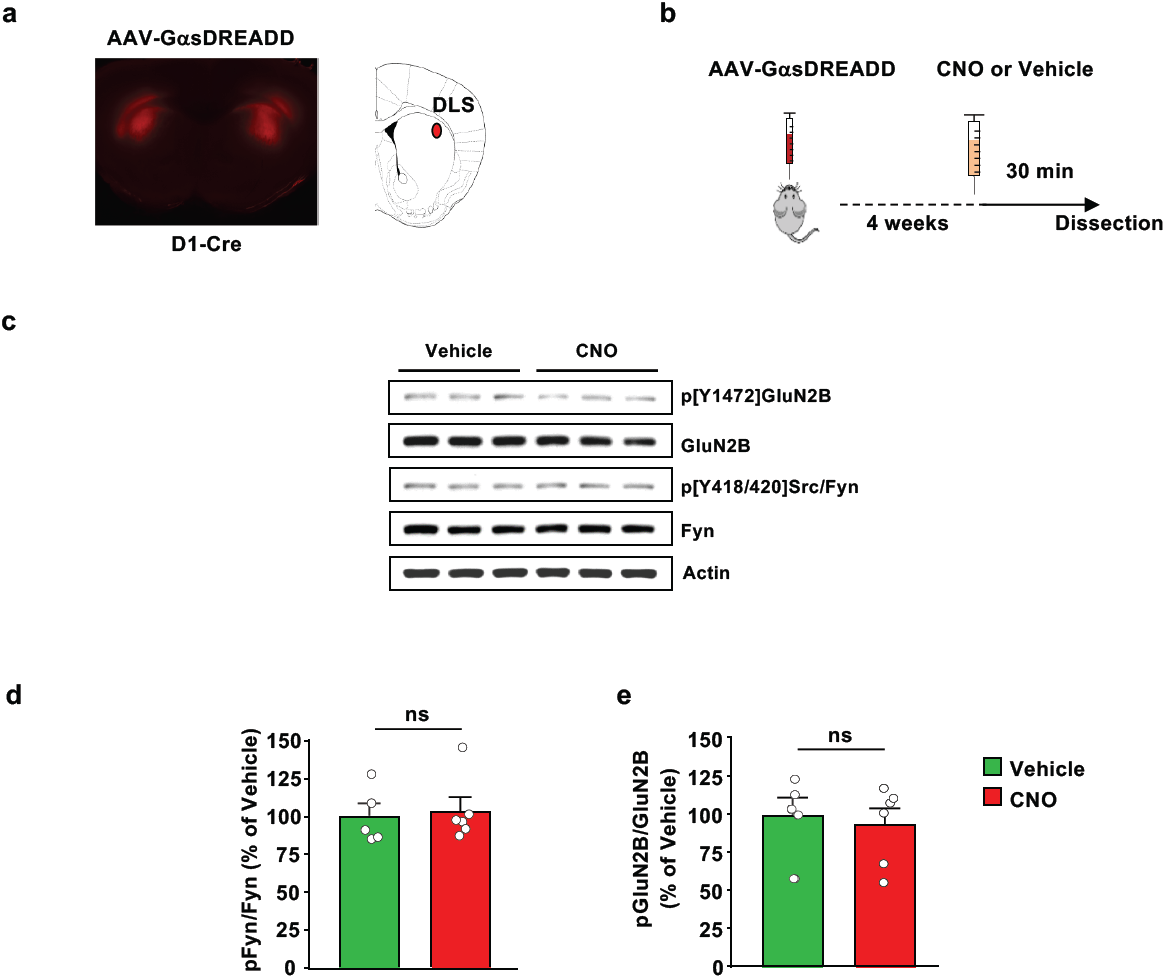
Stimulation of GαsDREADD in DLS dMSNs does not activate Fyn/GluN2B signaling. **(a)** The DLS of D1-Cre mice was infected bilaterally with AAV-DIO-rM3D(Gs)-mCherry (10^13^ vg/ml, 1 μl per side), and infection was evaluated 4 weeks later. Infection was visualized on EVOS FL microscope (Life Technologies, Carlsbad, CA), scale: 2x. (**b**) Timeline of experiment. Four weeks after surgery, vehicle (0.5% DMSO) or CNO (3 mg/kg) was systemically administered, and the DLS was dissected 30 minutes later. (**c**) Representative image of Fyn activation and GluN2B phosphorylation using anti-Tyr417/420[Src/Fyn] and anti-Tyr1472[GluN2B] antibodies, respectively. Total protein levels of Fyn, GluN2B and actin, which was used as a loading control, were measured in parallel. (**d-e**) Data are presented as the individual data points and mean densitometry values of the phosphorylated protein divided by the densitometry values of the total protein ± SEM and expressed as % of vehicle. Activation of GαsDREADD in DLS dMSNs does not change Fyn activation (**d**) (two-tailed unpaired t-test, t=0.2957, p=0.7742) and GluN2B phosphorylation (**e**) (two-tailed unpaired t-test, t=0.4105, p=0.6910). n=5-6 per treatment. ns: non-significant.

**Figure 4.**
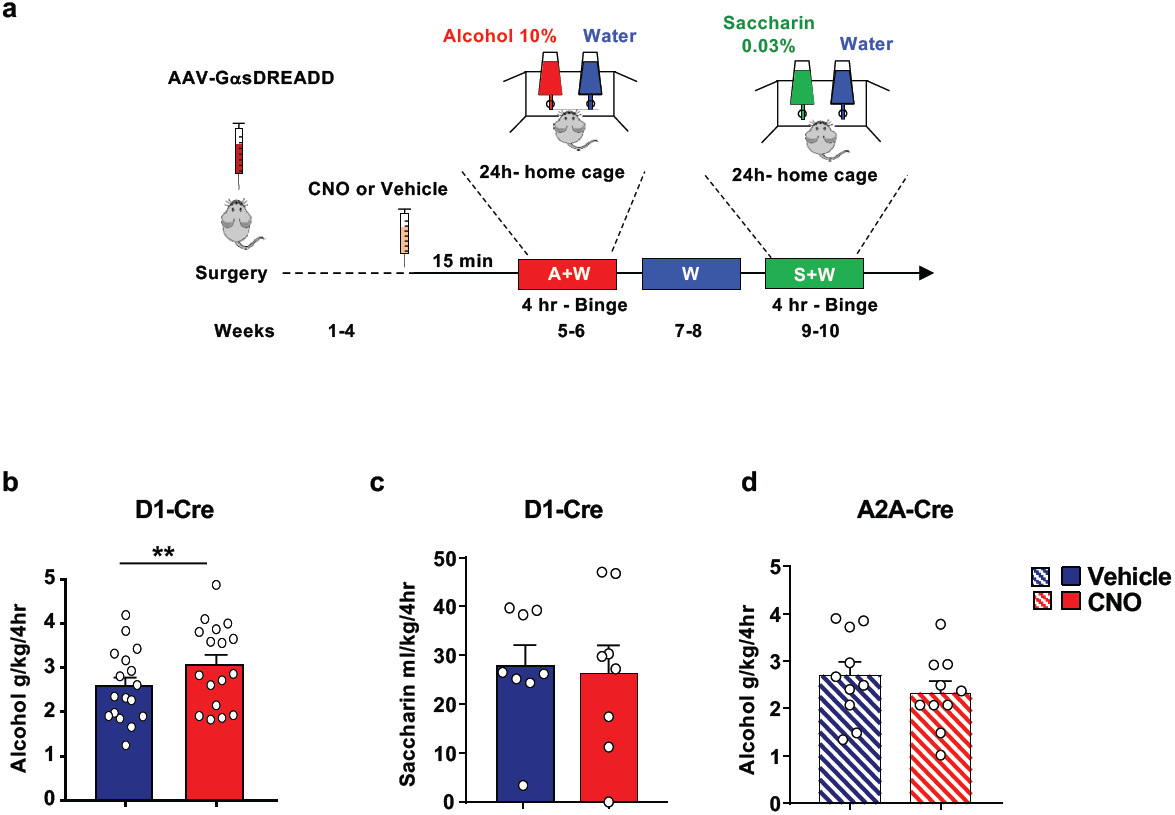
Stimulation of GαsDREADD in DMS dMSNs but not in iMSNs increases alcohol but not saccharine intake. (**a**) Timeline of experiment. The DMS of D1-Cre or A2A-Cre mice was infected bilaterally with AAV-DIO-rM3D(Gs)-mCherry (10^13^ vg/ml, 1 μl per side). (**b-e**) Four weeks after surgery, D1-Cre (**b**) or A2A-Cre (**d**) received a systemic administration of vehicle (0.5% DMSO) or CNO (3 mg/kg) 15 minutes before the beginning of the first 10% alcohol drinking session, and alcohol intake was measured 4 hours later. (**c**) At the completion of the alcohol drinking regimen, D1-Cre-mice had access to water only for two weeks. Afterwards, mice received a systemic administration of vehicle (0.5% DMSO) or CNO (3 mg/kg) 15 minutes before the beginning of a 0.03% saccharine drinking session, and saccharine and water intake were measured 4 hours later. (**b**) Activation of GαsDREADD in DMS dMSNs increases alcohol intake (two-tailed paired t-test, t=3.501, p=0.003). (**c**) Activation of GαsDREADD in DMS dMSNs does not alter saccharine intake (two-tailed paired t-test, t=0.2501, p=0.8097). (**d**) Activation of GαsDREADD in the DMS of A2A-Cre does not affect alcohol intake (two-tailed paired t-test, t=1.482, p=0.1724). Data are presented as individual data points and mean ± SEM. (**b**) n=17 per treatment, (**c**) n=10 per treatment, (**d**) n=8 per treatment. ** p<0.01.

A separate cohort of D1-Cre mice were infected with AAV-DIO-Gs-DREADD in the DMS. One month later, mice were systemically administered with vehicle (20% HPBCD) or AZD0530 (10 mg/kg) 3 hours prior to the beginning of the drinking session. Subsequently, animals received a systemic administration of vehicle (0.5% DMSO) or CNO, (3 mg/kg) 15 minutes before the beginning of the drinking session, and alcohol and water intake were measured 4 hours later (**Timeline, Figure 5a**). Mice were tested in a counterbalanced, within-subjects design.

**Figure 5.**
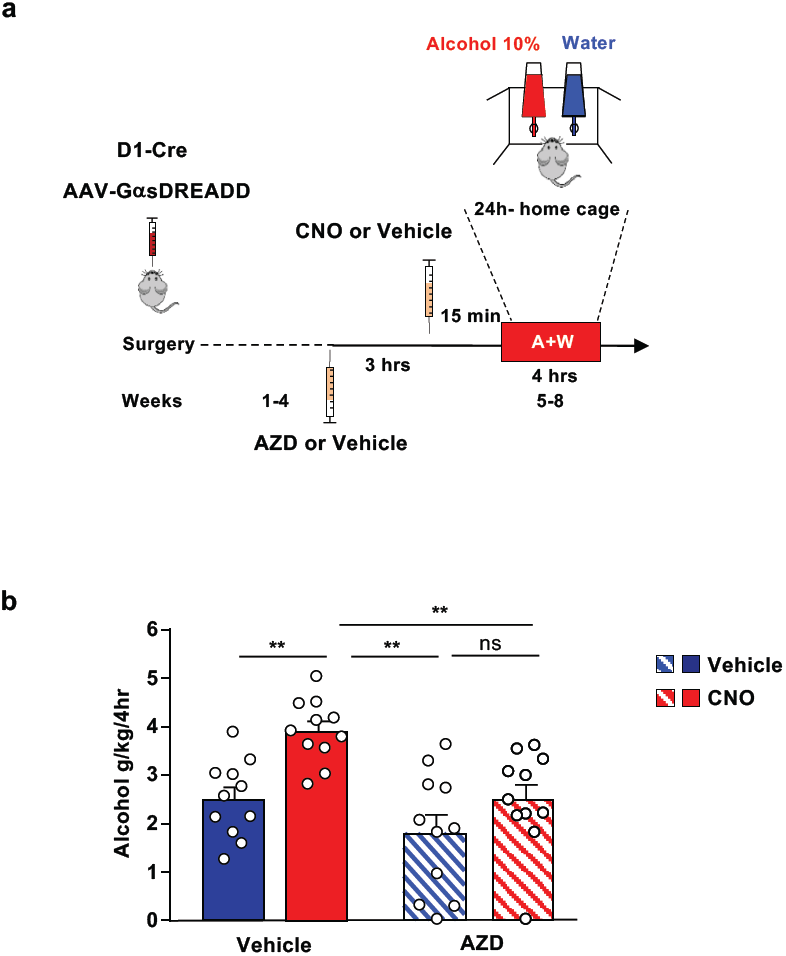
GαsDREADD-dependent increase in alcohol intake in DMS dMSNs requires Fyn. (**a**) Timeline of experiment. The DMS of D1-Cre mice was infected bilaterally with AAV-DIO-rM3D(Gs)-mCherry (10^13^ vg/ml, 1μl per side). Four weeks after surgery, animals received a systemic administration of vehicle (20% HPBCD) or AZD0530 (10 mg/kg) 3 hours before the beginning of a 10% alcohol drinking session. Subsequently, animals received a systemic administration of vehicle (0.5% DMSO) or CNO (3 mg/kg) 15 minutes before the beginning of the drinking session, and alcohol and water intake were measured 4 hours later. (**b**) AZD0530 blocks GαsDREADD-dependent enhancement of alcohol intake (Two-way RM ANOVA, effect of AZD0530: F_1,10_=9.177, p=0.0127; effect of CNO: F_1,10_=19.43, p=0.0013). Data are presented as individual values and mean ± SEM. n=11. ** p<0.01. ns: non-significant.

#### Two bottle choice - 0.03% Saccharin

D1-Cre/Ai14 mice underwent an intermittent saccharin intake procedure [45]. Specifically, 2 weeks after the end of the alcohol drinking paradigm during which mice consumed only water, mice had access to one bottle of water and one bottle of saccharin (0.03%) for one week (2 session) and saccharine and water intake were evaluated **(Timeline, Figure 1b).**

The DMS of D1-Cre mice were infected with AAV-DIO-Gs-DREADD. On test day, mice received a systemic administration of vehicle (0.5% DMSO) or CNO (3 mg/kg). Fifteen minutes later, mice had access to a bottle of saccharin (0.03%) and one bottle of water. Saccharin and water intake were measured 4 hours later (**Timeline, Figure 4a**). Mice were then given water only for one week and were tested again using a counterbalanced, within-subject design.

To measure the level of Fyn signaling activation by saccharine, C57BL/6 were subjected to one week of home cage 0.03% intermittent saccharine intake and the DMS was dissected at the last 4 hours drinking session (**Timeline, Supplementary Figure 1a**).

#### Fluid consumption measurements

Alcohol, saccharin and water bottles were presented in 50 ml graduated polypropylene cylinders with stainless-steel drinking spouts inserted through 2 grommets in front of the cage. Bottles were weighted before and 4 hours after the drinking session in order to determine the volume of consumed fluid. The weight of each mouse was measured the day before the drinking session to calculate the grams of alcohol intake per kilogram of body weight. The placement (right or left) of the bottles was alternated in each session to control for side preference. Two bottles containing water and alcohol in a cage without mice was used to evaluate the spillage due to the experimental manipulations during the test sessions. The spillage was always ≤0.2 ml. Alcohol (g/kg), saccharin (ml/kg) and water (ml/kg) intake were recorded at the end of each 4-hour drinking session.

### Statistical analysis

GraphPad Prism 7.0 (GraphPad Software, Inc., La Jolla, CA, USA) was used to plot and analyze the data. D’Agostino–Pearson normality test and *F*-test/Levene tests were used to verify the normal distribution of variables and the homogeneity of variance, respectively. Data were analyzed using the appropriate statistical test, including two-tailed unpaired or paired t-test, two-way analysis of variance (ANOVA) followed by post hoc tests as detailed in the Figure Legends. Experiments designed to evaluate the consequence of Fyn knockdown using FLEX-shFyn were analyzed with unpaired t-test. Experiments designed to test the contribution of cAMP signaling on alcohol intake using AAV-DIO-Gs-DREADD were analyzed with paired t-test since within subject design was used. All data are expressed as mean ± SEM, and statistical significance was set at *p*<0.05.

## Results

To determine whether Fyn’s actions are localized to DMS dMSNs and/or iMSNs, we used transgenic mice that express Cre recombinase and tdTomato specifically in dMSNs (D1-Cre/Ai14), or that express Cre recombinase and tdTomato specifically in iMSNs (A2A-Cre/Ai14), in combination with a Cre-dependent Flip Excision (FLEX) approach to downregulate *Fyn* mRNA in dMSNs or iMSNs, respectively, as described in [16]. First, we examined the consequence of Fyn knockdown in DMS dMSNs on alcohol drinking. To do so, the DMS of D1-Cre/Ai14 was infected bilaterally with a lentivirus expressing a short hairpin mRNA sequence targeting Fyn inserted in a FLEX cassette (FLEX-shFyn) (10^7^ pg/ml, 1.2 μl per site, two sites per hemisphere) or with a FLEX virus expressing a scramble sequence which was used as a control (FLEX-SCR, 10^7^ pg/ml, 1.2 μl per site, two sites per hemisphere) (**Figure 1a**). After 4 weeks, which enabled maximal viral infection and knockdown of the gene [16], mice were subjected to an additional week of IA20%2BC and alcohol intake was measured at the last 4 hours binge drinking session, a period in which mice drink the majority of alcohol, and in which mice reach a blood alcohol concentration (BAC) of over 80 mg% [43,46] (**Timeline, Figure 1b**). Alcohol intake was significantly reduced in D1-Cre/Ai14 mice infected with FLEX-shFyn in dMSNs as compared to the FLEX-SCR infected mice (**Figure 1c, Supplementary Table 1**).

We then assessed whether Fyn’s action in DMS dMSNs is specific for alcohol or is shared with other rewarding substances. To examine this question, we tested the consequences of Fyn knockdown in DMS dMSNs on the consumption of 0.03% saccharin (**Timeline Figure 1b**). Knockdown of Fyn in DMS dMSNs did not alter saccharin intake (**Figure 1d, Supplementary Table 1**) suggesting that Fyn is not activated by saccharine. As detailed in the introduction alcohol activates Fyn in the DMS of rodents [39,41,42] and that the activation of Fyn results in GluN2B phosphorylation [39,41,42]. To solidify the conclusion that Fyn in the DMS does not contribute to saccharine intake, we measured Fyn phosphorylation and GluN2B phosphorylation in the DMS of C57BL/6 mice consuming saccharine (Timeline, Supplementary Figure 1a). As shown in Supplementary Figure 1b-c, saccharine consumption (Supplementary Table 1), does not alter Fyn/GluN2B signaling in the DMS. Together, these results suggest that Fyn’s regulation of consummatory behavior in DMS dMSNs is specific for alcohol and is not generalized to other reinforcing agents.

Next, to determine if downregulation of Fyn in iMSNs also affects alcohol intake, the DMS of A2A-Cre/Ai14 mice was infected with FLEX-shFyn or FLEX-SCR. (**Timeline, Figure 1b**). After 4 weeks of recovery allowing maximal virus infection, mice were subjected to one week of IA20%2BC period, and alcohol intake was measured at the last 4 hours binge drinking session. We found that knockdown of Fyn in iMSNs does not alter alcohol intake (**Figure 1e, Supplementary Table 1**) suggesting that Fyn participates in mechanisms underlying alcohol consumption through its actions in dMSNs but not in iMSNs.

As stated above, D1Rs are selectively expressed in dMSNs [5]. D1Rs are coupled to Gαs/olf, and stimulation of Gαs/olf-coupled receptors activates cAMP/PKA signaling [6,7]. *Ex vivo* data suggest that Fyn is activated in the hippocampal neurons through the cAMP/PKA pathway [24,47-49]. **We therefore postulated that alcohol activates Fyn signaling which in turn facilitates alcohol drinking through the** stimulation of cAMP/PKA signaling in DMS dMSNs activates Fyn. To test this possibility, we utilized the Designer Receptor Exclusively Activated by Designer Drug (DREADD) system to remotely activate Gαs in DMS dMSNs or iMSNs [50]. First, the DMS of D1-Cre mice was infected bilaterally with AAV8-hSyn-DIO-rM3D(Gs)-mCherry (10^13^ vg/ml, 1 μl per hemisphere) (**Figure 2a**). Four weeks after surgery, vehicle or CNO (3 mg/kg) was administered systemically, and the DMS was harvested 30 minutes later (**Timeline, Figure 2b**). As shown in **Figure 2c-d**, CNO administration produced a robust increase in the phosphorylation and thus activation of Fyn in the DMS. To measure Fyn activation, we utilized anti-phosphoTyrosine417/420 antibodies which recognize the autophosphorylated active form of Fyn and Src [17,18]. To confirm that the kinase that was activated in response to CNO administration was indeed Fyn and not Src, we used another cohort of animals in which the DMS of D1-Cre mice was infected bilaterally with AAV8-hSyn-DIO-rM3D(Gs)-mCherry and treated with vehicle or CNS (3mg/kg), and conducted an immunoprecipitation assay in which Src was immunoprecipitated using specific anti-Src antibodies (Supplementary Figure 2a-b). The level of Src phosphorylation in response to CNO treatment was measured using the anti-phosphoTyrosine417/420 antibodies. As shown in Supplementary Figure 2c, Src was not activated upon stimulation of GαsDREADD in DMS dMSNs.

We also measured the phosphorylation level of the Fyn substrate, GluN2B [24], and found that Fyn activation in dMSNs was accompanied by the phosphorylation of GluN2B (**Figure 2c and e**). In contrast, CNO administration did not alter Fyn’s activity or GluN2B phosphorylation in the DLS, a neighboring striatal region which was not infected with AAV8-hSyn-DIO-rM3D(Gs)-mCherry (**Supplementary Figure 3**). Together, these data suggest that Fyn/GluN2B signaling is enhanced in response to remote activation of GαsDREADD in DMS dMSNs.

Next, we examined the level of Fyn signaling activation in the DLS dMSNs upon remote activation of GαsDREADD. To do so, the DLS of Drd1-Cre mice was infected bilaterally with AAV8-hSyn-DIO-rM3D(Gs)-mCherry (10^13^ vg/ml, 1 μl per hemisphere) (Figure 3a). Four weeks after surgery, vehicle or CNO (3 mg/kg) was administered systemically, and the DLS was harvested 30 minutes later (Timeline, Figure 3b). Strikingly, as shown in Figure 3c-e, remote activation of GαsDREADD in DLS dMSNs did not alter Fyn’s activity or GluN2B phosphorylation. These data suggest that GαsDREADD-dependent activation of Fyn signaling in dMSNs is centered in the DMS.

We then determined whether the cAMP-dependent activation of Fyn in dMSNs alters alcohol drinking. As Drd1-Cre mice consume large quantities of 20% alcohol ([51], **Figure 1c**), we used a lower alcohol concentration (10% v/v) in order to avoid a confounding ceiling effect of alcohol intake due to Gα*s*DREADD activation. The DMS of Drd1-Cre mice was infected bilaterally with AAV-hSyn-DIO-rM3D(Gs)-mCherry, four weeks later, vehicle or CNO (3 mg/kg) was administered systemically 15 minutes before the beginning of a 10% alcohol drinking session, and alcohol intake was measured after 4 hours (**Timeline, Figure 4a**). As shown in **Figure 4b, Supplementary Table 1**, remote activation of GαsDREADD in DMS dMSNs significantly increased alcohol intake.

Next, we examined whether remote activation of GαsDREADD in DMS dMSNs alters the consumption of saccharin. Two weeks after the end of the alcohol drinking experiment, vehicle or CNO (3 mg/kg) was administered systemically 15 minutes prior to the initiation of the saccharin (0.03%) drinking session, and saccharin intake was measured after 4 hours (**Timeline, Figure 4a**). Activation of GαsDREADD in dMSNs did not alter saccharin intake (**Figure 4c, Supplementary Table 1**) suggesting that the increase in consumption upon activation of GαsDREADD in DMS dMSNs is specific for alcohol.

We also examined whether remote activation of cAMP signaling in DMS iMSNs also alters alcohol intake. Interestingly, we found that CNO-dependent activation of GαsDREADD in DMS iMSNs does not affect alcohol intake (Figure 4d, Supplementary Table 1) suggesting that cAMP signaling in dMSNs but not iMSNs contributes to the development of excessive alcohol consumption.

Gomez et al. reported that CNO is converted to clozapine prior to binding and activating DREADDs [52], and furthermore, CNO was reported to produce some behavioral effects on its own [53,54]. Therefore, to ensure that the increase in alcohol intake was not due to off-target effects of the drug, we measured the level of alcohol consumption upon vehicle or CNO (3 mg/kg) treatment in D1-Cre mice that were not infected with AAV-hSyn-DIO-rM3D(Gs)-mCherry (**Timeline, Supplementary Figure 4a**). CNO administration did not alter alcohol intake in uninfected D1-Cre mice (**Supplementary Figure 4b, Supplementary Table 1)** suggesting that the increase in alcohol intake by CNO-dependent activation of Gα*s*DREADD in DMS dMSNs is not due to off target effects of the drug itself or its metabolite, clozapine.

Finally, we set out to test whether Fyn is required for GαsDREADD-dependent increase in alcohol consumption. To test this possibility, the DMS of D1-Cre mice was infected bilaterally with AAV-hSyn-DIO-rM3D(Gs)-mCherry, four weeks later, vehicle or the Src/Fyn inhibitor AZD0530 (10 mg/kg) [29,40,55] was administered 3 hours before the beginning of the 10% alcohol drinking session followed by the administration of vehicle or CNO (3 mg/kg) 15 minutes prior to the start of the session, and alcohol intake was measured after 4 hours (**Timeline, Figure 5a**). In accordance with **Figure 4b**, the amount of alcohol consumed by mice was elevated after the remote activation of GαsDREADD in DMS dMSNs, but the increase in alcohol intake was blocked when AZD0530 was administered prior to CNO (**Figure 4b, Supplementary Table 1**). In order to ensure that differences observed above are not due to a change in alcohol metabolism, we measured BAC after AZD0530 administration. To do so, mice received a systemic administration of vehicle or AZD0530 (10 mg/kg), 3 hour later, mice received a systemic administration of alcohol (2 g/kg), and BAC was measured 30 minutes later ((Timeline, Supplementary Figure 5a) As shown in Supplementary Figure 5b, BAC was similar in vehicle and AZD0530 treated mice suggesting that the drug does not affect alcohol metabolism. These results suggest alcohol intake is driven through the cAMP/PKA/Fyn signaling in DMS dMSNs.

## Discussion

Here, we present data to suggest that cAMP/PKA-dependent activation of Fyn kinase in the DMS dMSNs participates in mechanisms underlying the development of excessive alcohol intake. Specifically, we show that downregulation of Fyn levels in DMS dMSNs prior to the initiation of the IA20%2BC regimen attenuates alcohol intake. We further report that the stimulation of Gαs signaling in DMS dMSNs activates Fyn/GluN2B signaling and enhances alcohol intake in a Fyn-dependent manner. In contrast, knockdown of Fyn or activation of cAMP signaling in iMSNs does not alter alcohol intake and preference. Finally, we show that the cAMP/Fyn signaling in DMS dMSNs is specific for alcohol and does not contribute to mechanisms underlying consummatory behavior *per se*. Based on previous data showing that Fyn is activated in the DMS through the stimulation of D1R in DMS dMSNs [16], and since dopamine levels in the dorsal striatum are elevated by drugs of abuse including alcohol [56,57], we propose a model in which dopamine, released in the DMS in response to alcohol exposure, stimulates D1R/cAMP/PKA signaling in dMSNs which in turn activates Fyn to initiate neuroadaptations such as GluN2B phosphorylation that promote the development of excessive alcohol use (**Supplementary Figure 6**).

Our data suggest that Fyn in DMS dMSNs contributes to the development of **excessive** alcohol consumption. The DMS is essential for goal-directed behaviors [2,3], and dMSNs contribute to reward learning and reinforcement [58,59]. Thus, it is plausible that Fyn in dMSNs promotes reward learning which in turn initiates and maintains goal-directed alcohol seeking. This possibility is in line with the finding that systemic administration of the Src/Fyn inhibitor, AZD0530, attenuates goal-directed alcohol self-administration in mice [40]. Xie et al. previously showed that administration of the Src/Fyn inhibitor PP2 into the dorsal hippocampus attenuates context-dependent cocaine seeking [60], and more recently, Belin-Rauscent reported that oral administration of the Src/Fyn s inhibitor, Masitinib, attenuates self-administration of cocaine [61]. Thus, it would be of interest to determine whether Fyn in DMS dMSNs contributes to goal-directed seeking of other drugs of abuse. Furthermore, Goto et al. previously reported that mating behavior increases PKA activity in dorsal striatal dMSNs [62]. As PKA in DMS dMSNs is upstream of Fyn, it is plausible that the Fyn in these neurons plays a role in other goal-directed behaviors such as mating. Finally, Bocarsly et al. previously showed that enhanced D1R signaling in the dorsal striatum of mice is required for the consumption of alcohol despite negative consequences, and for the enhancement of alcohol-dependent hyperlocomotion [13]. Therefore, it plausible that Fyn in DMS dMSNs also contributes to other alcohol-dependent behaviors.

We found that stimulation of cAMP/PKA signaling activates Fyn but not Src in DMS dMSNs. This observation is in line with previous data showing that Fyn but not Src is activated in the dorsal striatum by alcohol exposure [38] or D1R stimulation [16]. However, it is plausible that the cAMP/PKA-dependent of Src signaling in other brain regions contribute to excessive alcohol use. For instance, Zhang et al. reported that opiate withdrawal activated Src in the locus coeruleus [63].

The data herein and previous findings [31] suggest that Fyn exerts its action on alcohol drinking in dMSNs through the phosphorylation and activation of GluN2B. However, we cannot exclude the possibility that other molecular transducers of Fyn contribute to the development of excessive alcohol use. For example, protein translation plays a critical role in mechanisms underlying AUD [64], and Fyn was shown to enhance protein translation in oligodendrocytes [65] and in neurons [66]. Fyn was also shown to promote ERK1/2 phosphorylation in oligodendrocytes [67], and NfkB signaling in microglia [68]; both signaling cascades have been linked to alcohol use [30,69]. Finally, Fyn was reported to phosphorylate the metabotropic glutamate receptor 1 (mGluR1) [70] and the Collapsin Response Mediator Protein 2 (CRMP2) [71], which have also been implicated in alcohol’s actions in the brain [72,73]. Exploring the contribution of these substrates and others in DMS dMSNs to the neuroadaptations underlying AUD merit further investigation.

It is highly likely that the activation of PKA in DMS dMSNs produces additional cellular consequences. For example, PKA phosphorylates the Striatal-Enriched Protein Tyrosine Phosphatase (STEP) [49] resulting in the inhibition of the activity of the phosphatase [49]. STEP is an endogenous Fyn inhibitor and is responsible for the termination of Fyn activation [49]. We previously reported that alcohol increases PKA phosphorylation of STEP in the DMS, and that knockdown of STEP in the DMS [41], or global knockout of the phosphatase increases alcohol intake [74]. It is therefore plausible that one of the consequences of PKA activation in DMS dMSNs is the phosphorylation of STEP, enabling Fyn in these neurons to stay active for a prolonged period of time.

Using the DREADD/CNO methodology, we report that the activation of GαsDREADD in dMSNs but not in iMSNs initiates the consumption of alcohol. We further show that GαsDREADD-dependent increase in alcohol intake dependent, at least in part, on Fyn. To our knowledge this is the first study that provides a link between GαsDREADD activation and alcohol consummatory behavior. Cheng et al. previously showed that GαiDREADD-mediated inhibition of iMSNs enhances alcohol intake [75]. Stimulation of Gαs-coupled receptors activates adenylate cyclase that increases cAMP production, whereas the stimulation of Gαi-coupled receptors inhibits adenylate cyclase activity and cAMP production [7]. Thus, it is plausible that the development of excessive alcohol consumption depends on the inhibition of cAMP signaling in iMSNs and on the activation of cAMP signaling in dMSNs.

Our data suggest that alcohol-dependent molecular adaptations are highly specific and are segregated to a subpopulation of neurons. Future studies are necessary to determine whether this cell-type specificity is unique for the cAMP/PKA/Fyn signaling or that this is a common feature shared by some or all of the molecular targets of alcohol [30].

Although we provide a strong evidence linking PKA signaling in dMSNs to alcohol drinking behaviors, we cannot exclude the possibility that other cAMP effectors such as the guanine nucleotide exchange factor EPAC (exchange protein directly activated by cAMP) [76], and/or cyclic nucleotide-gated ion channels (CNGC) [77] also contribute to the development of excessive alcohol use. Finally, further research is required to monitor cAMP production and PKA activation in behaving animals. Recent advances in the development of reporters for cAMP [78], and PKA [79] will enable a spatial and temporal analysis of cAMP/PKA signaling in animals consuming alcohol.

Finally, we previously showed that treatment of mice with AZD530 attenuates alcohol-dependent Fyn activation and GluN2B phosphorylation in the DMS [40], and reduces goal-directed alcohol seeking. We show herein that the enhancement of alcohol intake upon activation of GαsDREADD in DMS dMSNs is inhibited upon the administration of AZD530. AZD530 is well-tolerated in humans, as phase I and II clinical trials indicate that the drug does not produce significant side effects [80,81] and our preclinical mouse studies show that systemic administration of the drug does not alter basal levels of locomotion [40], and does not change BAC. Together, these data give rise to the potential use of AZD530 in alcohol use disorder.

## Acknowledgements

We thank AstraZeneca for providing us with AZD5030. The authors thank Ellanor Whiteley for her contribution.

## Author Contributions

Y.E contributed to the design of the experiments, the acquisition of data, data analysis, and to the preparation and revision of the manuscript. N.M, S.A.S and K.P contributed to the design of the experiments, the acquisition of the data and data analysis. M.F.A. and D.S contributed to the acquisition of data. V.A.A contributed to the conception of the study. D.R. contributed to the conception of the study, the design of the experiments and wrote the manuscript.

## Supplementary Information

### Supplementary Materials and Methods

#### Reagents

Goat anti-GluN2B antibodies (Lot # C0606) and rabbit anti-Fyn antibodies (Lot # E152) and mouse IgG were purchased from Santa Cruz Biotechnology (Santa Cruz, CA). Rabbit anti-[pY1472]GluN2B antibodies (Lot # 3) and rabbit anti-[pY418/420]Src/Fyn antibodies (Lot# 21) were purchased from Cell Signaling Technology (Beverly, MA). Mouse anti-Actin antibodies (Lot # 127M4866V), Clozapine N-Oxide (CNO), and Phosphatase inhibitor cocktails 2 and 3 were purchased from Sigma Aldrich (St. Louis, MO). Donkey anti-rabbit horseradish peroxidase (HRP), donkey anti-goat horseradish peroxidase (HRP) and donkey anti-mouse horseradish peroxidase (HRP) conjugated secondary antibodies were purchased from Jackson ImmunoResearch (West Grove, PA). Enhanced chemiluminescence (ECL) was purchased from GE Healthcare (Marlborough, MA). Agfa X-Ray Film was purchased from VWR (Radnor, PA). EDTA-free protease inhibitor cocktail was purchased from Roche (Indianapolis, IN). Bicinchoninic acid (BCA)™ protein assay kit was purchased from Pierce (Rockford, IL). Nitrocellulose membrane and mouse anti-Src antibodies was purchased from EMD Millipore (Billerica, MA). Protein G was purchased from Life Technologies (Carlsbad CA). Adeno-Associated Virus (AAV) AAV-hSyn-DIO-rM3D(Gs)-mCherry (AAV vector serotype 5) was produced from and purified by the Viral Vector Core Facility at Duke University. HIV-1 p24 antigen ELISA Kit was purchased from ZeptoMetrix (Buffalo, NY). AZD5030 was a generous gift from AstraZeneca.

#### Collection of brain samples

Animals were euthanized and brains were rapidly dissected on ice using a 1 mm brain block. The DMS and the dorsolateral striatum (DLS) were isolated from a 1 mm thick coronal section located between +1.18 mm and +0.74 mm anterior to bregma, −2.45 to −3.75 mm below the brain surface and +/-0.75 to 1.25 mm (DMS) and +/- 1.75 to 2.5 mm (DLS) medial to the midline according to the Franklin and Paxinos stereotaxic atlas (3^rd^ edition). Tissues were dissected on an anodized aluminum block on ice, collected into 1.5 ml Eppendorf tubes, and immediately homogenized in 300 μl RadioImmuno Precipitation Assay (RIPA) buffer containing (in mM: 50 Tris-HCl, pH 7.6, 150 NaCl, 2 EDTA), and 1% NP-40, 0.1% SDS and 0.5% sodium deoxycholate and protease and phosphatase inhibitor cocktails. Samples were homogenized by a sonic dismembrator. Protein content was determined using a BCA kit.

#### Western Blot analysis

Equal amounts of homogenates from individual mice (30 μg) were resolved on NuPAGE Bis-Tris gels and transferred onto nitrocellulose membranes. Blots were blocked in 5% milk-PBS, 0.1% Tween 20 for 30 minutes and then incubated overnight at 4°C with anti-[pY418/420Src/]Fyn (1:500) or anti-[pY1472]GluN2B (1:250) antibodies. Membranes were then washed and incubated with HRP-conjugated secondary antibodies for 2 hours at room temperature. Bands were visualized using ECL. Afterwards, membranes were stripped for 30 minutes at room temperature in a buffer containing 25 mM Glycine-HCL and 1% (w/v) SDS, pH 3.0, reprobed with anti-Fyn (1:500), anti-GluN2B (1:500), and anti-Actin (1:5000) antibodies and processed as described above. Optical density of the relevant band was quantified using ImageJ 1.44c software (NIH).

#### Immunoprecipitation

Immunoprecipitation assay was conducted as described in [1]. Brain regions were dissected as described above, and tissues were then homogenized in immunopreci,pitation (IP) buffer containing (in mM: 150 NaCl, 10 Tris-HCl pH 7.4, 1 EDTA, 1 EGTA) 1% Triton-X, protease and phosphatase inhibitor cocktails. Homogenates were pre-cleared by incubation with protein G agarose for 1 hour at 4°C. Samples were then spun down and supernatant protein quantity was determined using BCA protein assay. IPs were performed by combining 1 μg of the appropriate antibody with 500 μg lysate diluted in IP buffer to a total volume of 1 ml. Following overnight incubation at 4°C, protein G agarose was added, and the mixture was incubated at 4°C for 4 hours. Protein G was washed extensively with IP buffer and pellets were re-suspended in 40 μl of 2x Laemmli buffer and incubated at 95°C for 10 minutes. The proteins in the supernatant were separated by SDS-PAGE gels and visualized by ECL.

#### Blood Alcohol Concentration Measurement

Blood alcohol concentration (BAC) procedure was conducted as described in [2] with modifications. Blood was collected intracardially in heparinized capillary tubes from C57BL/6 mice 30 minutes after an i.p. injection of 2 g/kg of alcohol. Serum was extracted with 3.4% trichloroacetic acid followed by a 5-minute centrifugation at 420 g and assayed for alcohol content using the NAD-NADH enzyme spectrophotometric method [3,4]. BACs were determined by using a standard calibration curve.

#### Preparation of FLEX-shRNA-Fyn and FLEX-SCR

Details regarding the construction, preparation and characterization of lenti-virus expressing *FLEX-shRNA-Fyn* and *FLEX-SCR* are described in [1]. Virus titer was determined using HIV-1 p24 antigen ELISA Kit per the manufacturer’s instruction.

#### Preparation of solutions

Alcohol solution was prepared from absolute anhydrous alcohol (190 proof) diluted to 10% or 20% alcohol (v/v) in tap water. Saccharin solution was diluted to 0.03% sucrose (v/v) in tap water. CNO was dissolved in DMSO and then diluted in saline to a concentration of 3 mg/kg in 0.5% DMSO. Vehicle consisted of 0.5% DMSO in saline solution. AZD5030 was dissolved in 20% hydroxypropul beta cyclodexrin (HPbCD) diluted in saline and administered at a concentration of 10 mg/kg [5,6]. Vehicle consisted of 20% HPbCD diluted in saline.

## Supplementary Table and Figure Legends

**Supplementary Table 1.**
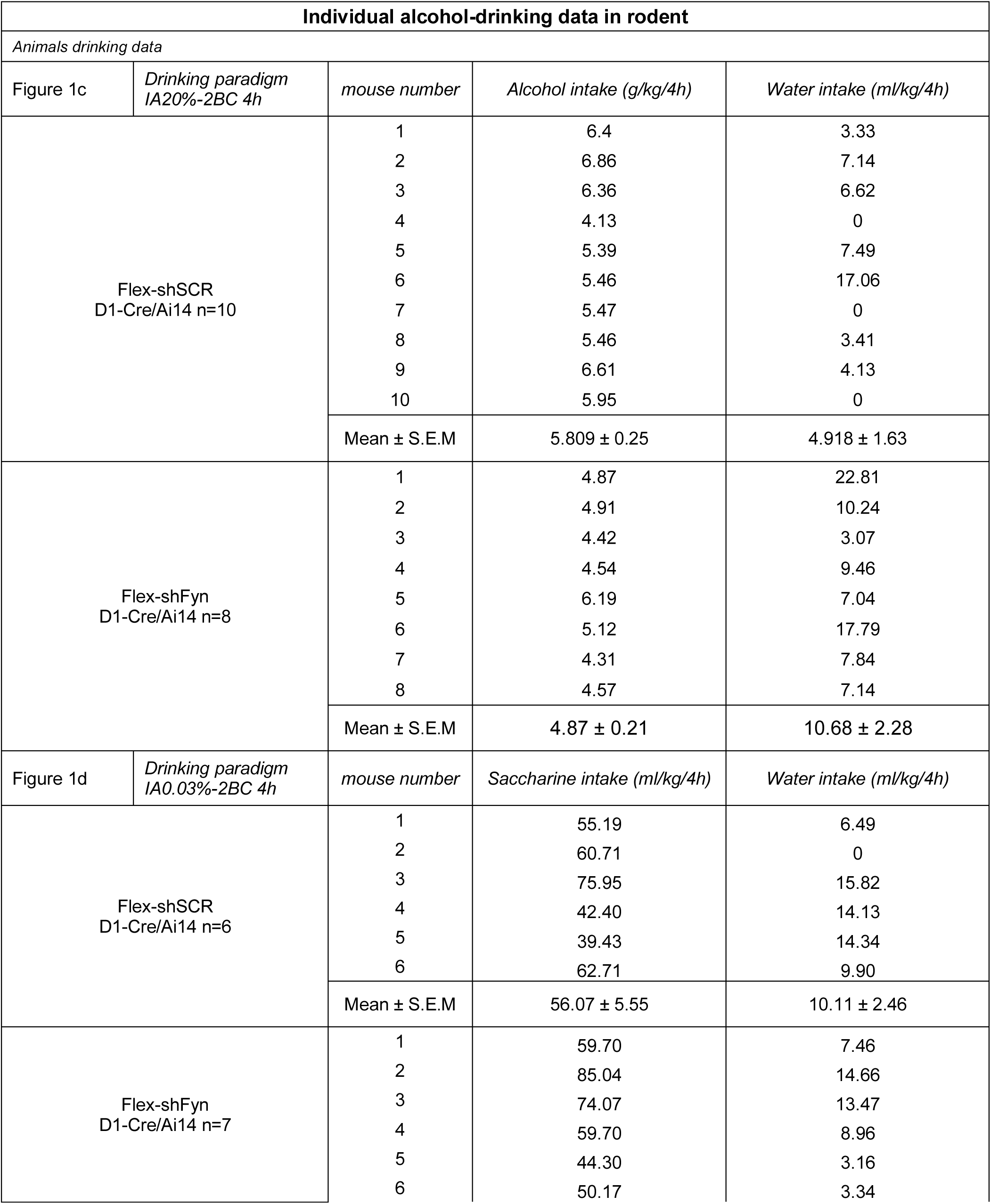
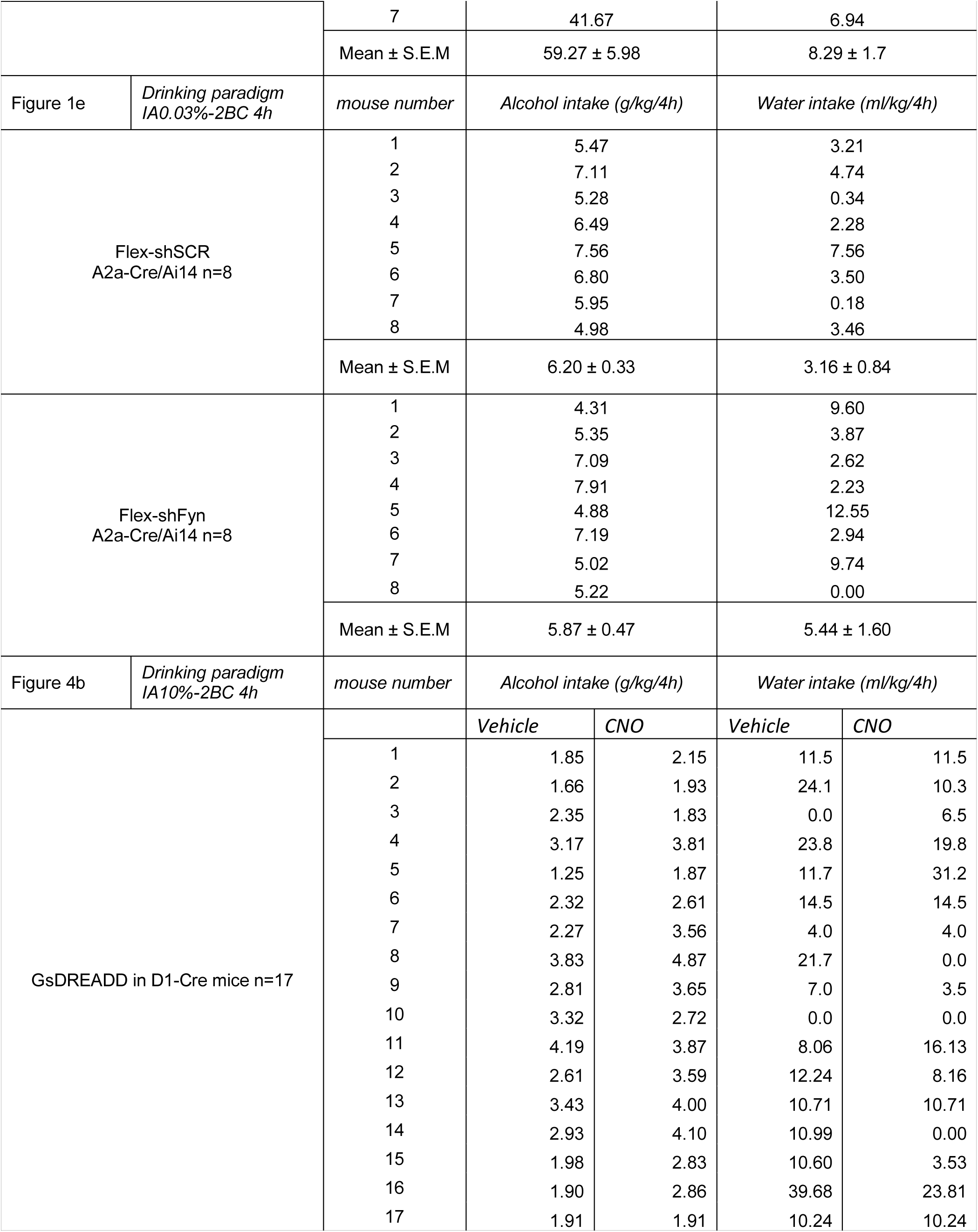
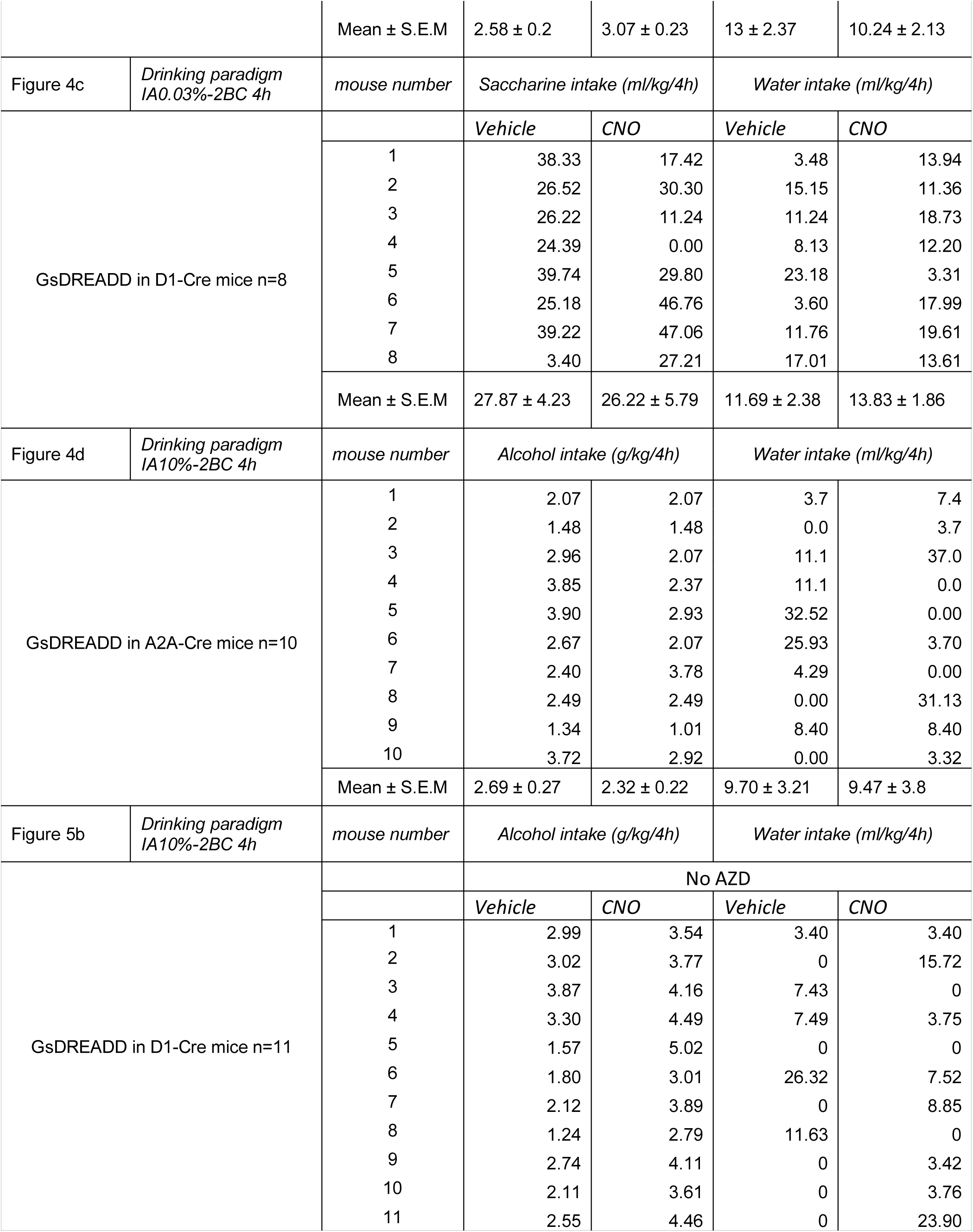
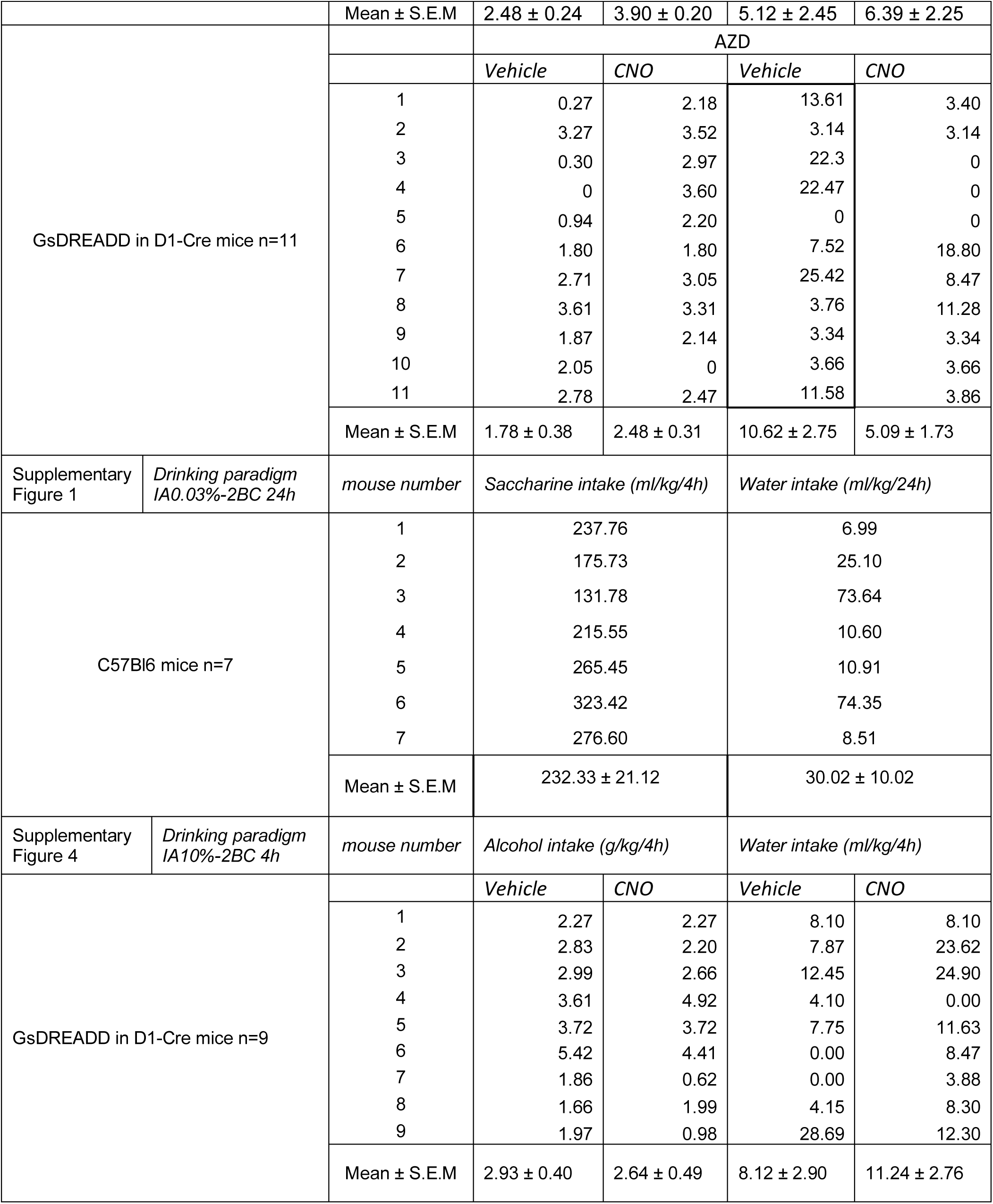
Individual alcohol and saccharin drinking data. Alcohol (g/kg) and saccharin (ml/kg) intake data from individual animals. The mean of the consumed amount is reported, labeled according to experiments (10% alcohol, 20% alcohol or 0.03% saccharine) and experimental manipulation.

**Supplementary Figure 1.**
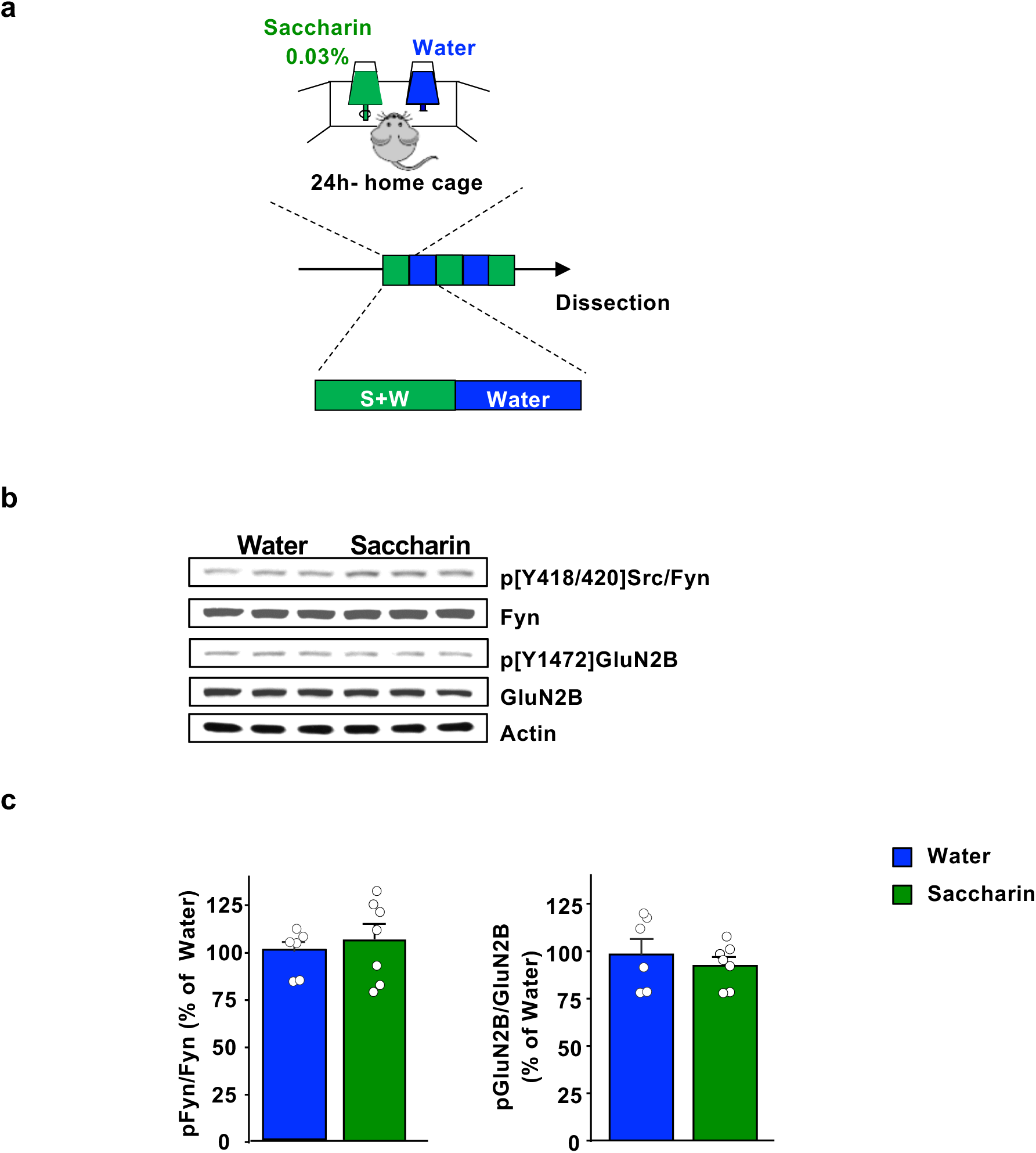
Saccharin drinking does not activate Fyn/GluN2B signaling in the DMS. Timeline of experiment. C57BL/6 mice underwent one week of home cage intermittent access to 0.03% saccharin. After the last drinking session, the DMS was dissected and the phosphorylation of Fyn and GluN2B were measured by Western blot analysis using anti-Tyr417/420[Src/Fyn] and anti-Tyr1472[GluN2B] antibodies, respectively. Total protein levels of Fyn, GluN2B and actin, which was used as a loading control, were measured in parallel. **(b)** Representative image of Fyn and GluN2B phosphorylation in the DMS. **(c)** Saccharin drinking does not increase Fyn activation (two-tailed unpaired t-test, t=0.6438, p=0.5329) or GluN2B phosphorylation (two-tailed unpaired t-test, t=0.7715, p=0.4567) in the DMS. Data are presented as the individual data points and mean densitometry values of the phosphorylated protein divided by the densitometry values of the total protein ± SEM and expressed as % of water. n=6-7 per treatment.

**Supplementary Figure 2.**
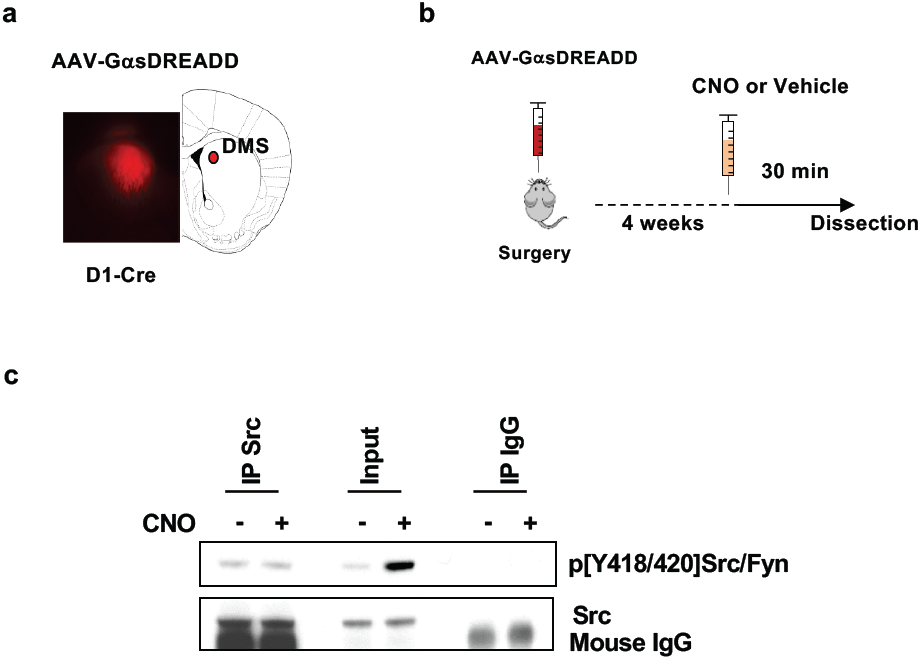
Stimulation of GαsDREADD in the DMS dMSNs does not activate Src. **(a)** The DMS of D1-Cre mice was infected bilaterally with AAV-DIO-rM3D(Gs)-mCherry (10^13^ vg/ml, 1 μl per side), and infection was evaluated 4 weeks later, scale: 2x. (b) Timeline of experiment. Four weeks after surgery, vehicle (0.5% DMSO) or CNO (3 mg/kg) was systemically administered, and the DMS was dissected 30 minutes later. Src was immunoprecipitated using anti-Src antibodies **(c)**. Mouse IgG was used as a negative control (IP IgG). Src immunoreactivity (lower panel) as well as Tyr417/420[Src/Fyn] phosphorylation (upper panel) were determined by westernblot analysis. Protein levels in the total lysates were measured in parallel (Input). n = 6 mice per treatment.

**Supplementary Figure 3.**
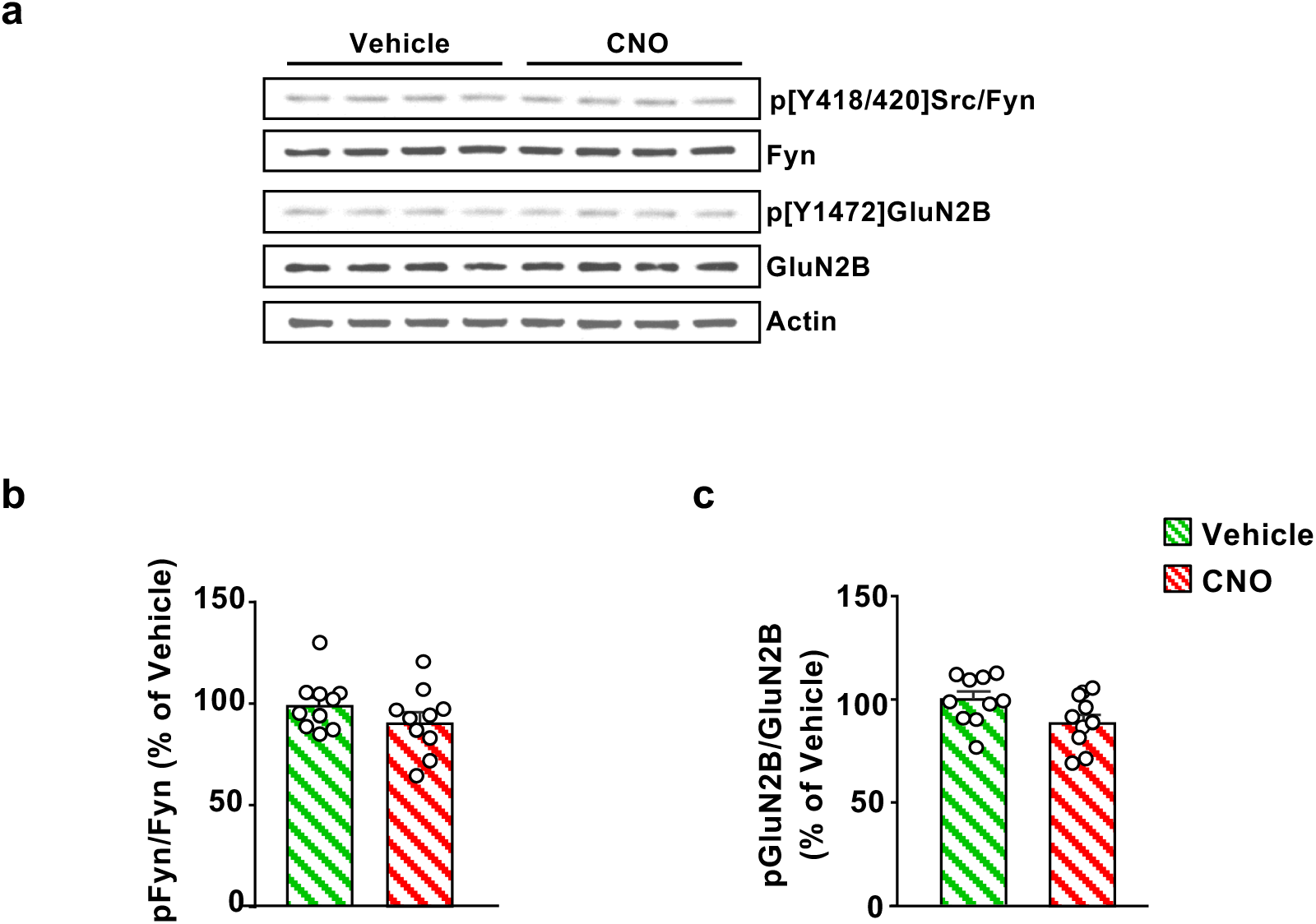
Stimulation of GαsDREADD in the DMS dMSNs does not activate Fyn/GluN2B in the DLS. The DLS was dissected from the same cohort of animals depicted in Figure 2. Fyn and GluN2B phosphorylation were measured by Western blot analysis using anti-Tyr417/420[Src/Fyn] and anti-Tyr1472[GluN2B] antibodies, respectively. Total protein levels of Fyn, GluN2B and actin, which was used as a loading control, were measured in parallel. (**a**) Representative image of Fyn and GluN2B phosphorylation in the DLS. (**b**,**c**) Activation of GαsDREADD in DMS dMSNs does not increase Fyn activation (**b**) (two-tailed unpaired t-test, t=1.262, p=0.2239), or GluN2B phosphorylation (**c**) (two-tailed unpaired t-test, t=1.838, p=0.0836) in the DLS. Data are presented as the individual data points and mean densitometry values of the phosphorylated protein divided by the densitometry values of the total protein ± SEM and expressed as % of vehicle. n=10 per treatment.

**Supplementary Figure 4.**
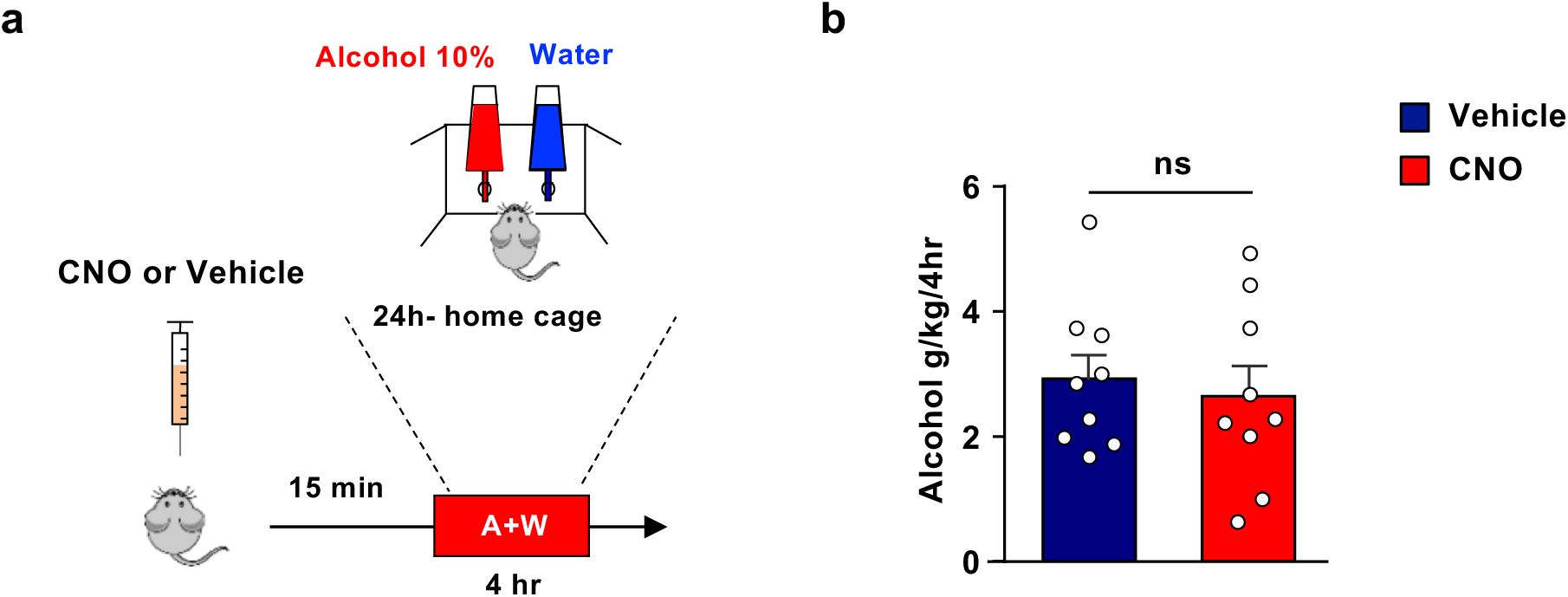
Administration of CNO in D1-Cre mice does not alter the consumption of 10% alcohol. (**a**) Timeline of experiment. D1-Cre mice received a systemic administration of vehicle (0.5% DMSO) or CNO (3 mg/kg) 15 minutes before the beginning of a 10% alcohol drinking session, and alcohol and water intake were measured 4 hours later. Mice were then given water for one week and were tested again using a counterbalanced within subject design. Data are presented as individual values and mean ± SEM. (**b**) CNO administration does not alter 10% alcohol intake (two-tailed paired t-test, t=1.063, p=0.3187). n=9.

**Supplementary Figure 5.**
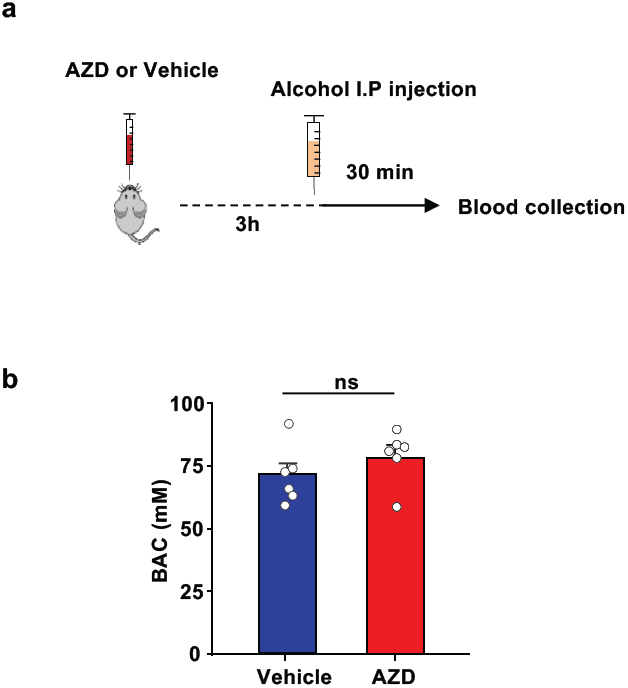
Systemic administration of AZD0530 does not affect blood alcohol concentration. **(a)** Timeline of experiment. C57BL/6 mice received a systemic administration of vehicle or AZD0530 (10 mg/kg). Three hour later, mice received a systemic administration of alcohol (2 g/kg), and BAC was measured 30 minutes later. **(b)** BAC is unaltered in AZD treated mice (unpaired t-test, t=1.216, p=0.2519). Data are presented as mean ± SEM. n = 6 per group.

**Supplementary Figure 6.**
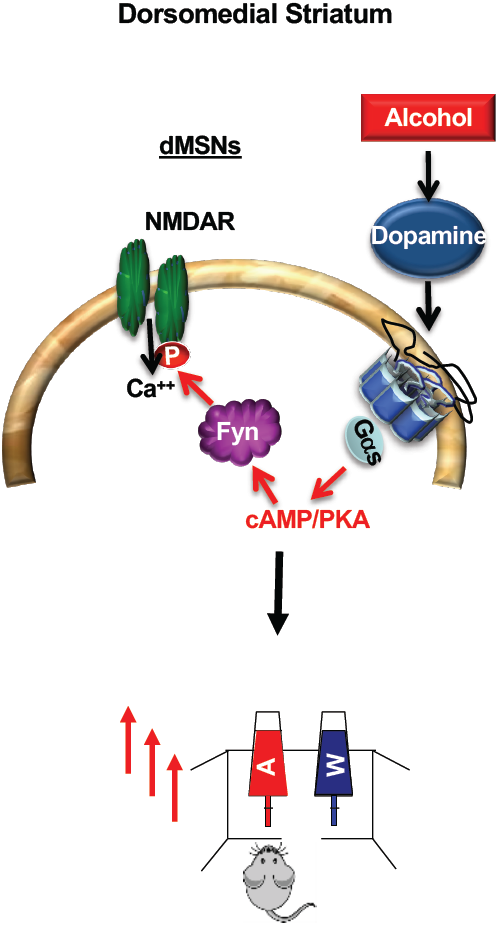
Activation of Fyn signaling in dMSNs promotes alcohol drinking. Cartoon depicting a model in which alcohol increases dopamine levels in the DMS resulting in the activation of the dopamine D1Rs in dMSNs. Dopamine binding to D1R stimulates cAMP/PKA signaling, which in turn activates Fyn. Fyn phosphorylates GluN2B resulting in the enhancement of GluN2B activity. These molecular adaptations contribute to the reinforcement of alcohol drinking behavior.

